# Host-pathogen coevolution promotes the evolution of general, broad-spectrum resistance and reduces foreign pathogen spillover risk

**DOI:** 10.1101/2023.08.04.548430

**Authors:** Samuel V. Hulse, Janis Antonovics, Michael E. Hood, Emily L. Bruns

## Abstract

Genetic variation for disease resistance within host populations can strongly impact the spread of endemic pathogens. In plants, recent work has shown that within-population variation in resistance can also affect the transmission of foreign spillover pathogens if that resistance is general. However, most hosts also possess specific resistance mechanisms that provide strong defenses against coevolved endemic pathogens. Here we use a modeling approach to ask how antagonistic coevolution between hosts and their endemic pathogen at the specific resistance locus can affect the frequency of general resistance, and therefore a host’s vulnerability to foreign pathogens. We develop a two-locus model with variable recombination that incorporates both general (resistance to all pathogens) and specific (resistance to endemic pathogens only). We find that introducing coevolution into our model greatly expands the regions where general resistance can evolve, decreasing the risk of foreign pathogen invasion. Furthermore, coevolution greatly expands which conditions maintain polymorphisms at both resistance loci, thereby driving greater genetic diversity within host populations. This genetic diversity often leads to positive correlations between host resistance to foreign and endemic pathogens, similar to those observed in natural populations. However, if resistance loci become linked, the resistance correlations can shift to negative. If we include a third, linkage modifying locus into our model, we find that selection often favors complete linkage. Our model demonstrates how coevolutionary dynamics with an endemic pathogen can mold the resistance structure of host populations in ways that affect its susceptibility to foreign pathogen spillovers, and that the nature of these outcomes depends on resistance costs, as well as the degree of linkage between resistance genes.

## Introduction

Natural populations are frequently confronted with foreign pathogen spillovers, where a host becomes infected with a novel pathogen (Borremans et al., 2019). Warming climates and habitat destruction are driving changes in species distributions, which can increase spillover risks via increased pathogen exposure rates (Jones et al., 2008). Understanding the drivers of susceptibility to foreign pathogens is therefore essential. At the species level, the phylogenetic distance between naïve and ancestral hosts is a key predictor of host shift potential, where pathogens typically are more infectious against closely related hosts (Parker et al., 2017; Streicker et al., 2010). However, within species, populations often harbor genetic variation in disease resistance (Baker & Antonovics, 2012; Laine, 2004; Thrall et al., 2001), including variation in resistance to diseases with which they share no coevolutionary history (Antonovics et al., 2002; Best & Kerr, 2000; Savage & Zamudio, 2011). Processes that drive the evolution of higher host resistance to endemic pathogens could therefore increases resistance to foreign pathogens if that resistance is positively transitive, that is, resistance to endemic pathogens is positively correlated with resistance to foreign pathogens.

Recently, Lerner et al., (2021) found significant positive correlation between genetic resistance to an endemic fungal pathogen and to a related foreign pathogen in the herbaceous plant *Silene vulgaris*. Based on these results, the authors developed a model that assumed resistance transitivity was determined by a single pleiotropic locus and found that the slope of the resistance correlation can affect the risk of foreign pathogen invasions. However, we do not yet understand the evolutionary forces that could influence the slope of the resistance correlation or maintain genetic diversity for resistance to foreign pathogens. One potential clue lies in the presence of a few notable and repeatable outliers in Lerner’s data, where some host genotypes were more susceptible to the foreign pathogen than to the endemic pathogen. These outliers suggest some genotypes may have specific resistance against the endemic pathogen but lack more general forms of resistance which could protect against foreign pathogens.

Plants and animals have evolved general forms of resistance that are effective against a wide range of pathogens, which stimulate responses such as inflammation in animals or, thickening of cell walls and production of anti-microbial peptides in plants (Nürnberger et al., 2004). Since these mechanisms are broadly effective, general resistance should protect hosts against foreign pathogens. However, hosts have also evolved highly specific forms of resistance which are triggered by a single species of endemic pathogen (or often even a specific pathogen genotype) (Märkle et al., 2022). Previously, using an adaptive dynamics approach, we found that strong selection from a single, invariant endemic pathogen can favor the evolution of specific resistance and drive the loss of general resistance, leaving populations more vulnerable to foreign pathogens (Hulse et al., 2023). However, our prior model did not account for host-pathogen coevolution.

Specific resistance is particularly vulnerable to host-pathogen coevolution because it generates strong positive selection for pathogens genotypes that can evade this resistance. In plants, this dynamic has been formalized with the gene-for-gene model (Flor, 1956). In the gene-for-gene model, a specific resistance gene (*R* gene) recognizes a particular effector molecule secreted by a pathogen (coded by an *Avr* gene). Mutations in the *Avr* gene that avoid recognition by the *R* gene (“virulent” form) allow the pathogen to successfully infect resistant hosts, often driving cyclical dynamics in host resistance and pathogen virulence (Tellier & Brown, 2007). Many natural systems have been shown to exhibit temporally and spatially variable selection for resistance due to gene-for-gene dynamics (Laine, 2005; Thrall et al., 2012; Thrall & Burdon, 2003). In agriculture, the introduction of new *R* genes is reliably followed by the rapid evolution of corresponding pathogen virulence (Carson, 2011; Miller et al., 2020). In contrast, general forms of resistance in crop species are often considered to be more “durable” over time, and do not generate the same rapid coevolutionary response from pathogens (Mundt, 2014; Poland et al., 2009). However, few studies have investigated how coevolutionary dynamics at specific resistance loci affects the evolution of general resistance (Koskella et al., 2011; Koskella & Parr, 2015).

Here, we develop a 2-locus host-pathogen model to investigate the joint evolution of general resistance (affecting both endemic and foreign pathogens) and specific resistance (affecting only endemic pathogens). We first ask how the introduction of a pathogen that can evolve in response to specific, but not general resistance shapes the evolution of both forms of resistance. For models to be relevant to natural systems, evolutionary outcomes must be described in terms measurable for empiricists. Therefore, we use our model to estimate the family-level correlation in resistance (transitivity) to endemic and foreign pathogens, a measure easily quantifiable in natural populations, where the exact loci underlying resistance are often unknown. Our results suggest that the degree of recombination between general and specific resistance strongly affects evolutionary outcomes. We therefore introduced a third locus that modifies the recombination rate, to determine whether selection favors the evolution of linkage between general and specific resistance. We find that coevolution expands the range of conditions under which general resistance can evolve, and can drive both positive and negative resistance correlations, depending on resistance costs and genetic linkage between resistance loci.

## Methods

We develop a compartmental model (Eq 1-2 below) that is loosely based on the dynamics of the anther-smut disease (caused by *Microbotryum silenes-inflatae*) on the herbaceous plant *Silene vulgaris*. In this system, transmission is frequency-dependent, and infection causes sterilization without significant effects on mortality (Thrall et al., 1995). For simplicity, we assume haploid hosts, with a locus for general resistance and a locus for specific resistance. General resistance gives hosts a moderate level of resistance against all pathogen genotypes and species, while specific resistance gives hosts a high level of resistance against a single endemic pathogen genotype. Each locus has an allele (*g* and *s*) which confer no resistance benefits and no costs, as well as an allele (*G* and *S*) which confer both resistance benefits and fecundity costs, giving our model four possible host genotypes (Table 1).

**Table 1.** Resistance levels and birthrates for each genotype.

| Host genotype | Host fecundity | Avirulent endemic transmission | Virulent endemic pathogen transmission | Foreign pathogen transmission |
| --- | --- | --- | --- | --- |
| *GS* | $(1 - c_G)(1 - c_S)$ | $\beta(1 - r_G)(1 - r_S)$ | $\beta(1 - r_G)$ | $\beta(1 - r_G)(1 - r_f)$ |
| *Gs* | $1 - c_G$ | $\beta(1 - r_G)$ | $\beta(1 - r_G)(1 - r_v)$ | $\beta(1 - r_G)(1 - r_f)$ |
| *gS* | $1 - c_S$ | $\beta(1 - r_S)$ | $\beta$ | $\beta(1 - r_f)$ |
| *gs* | 1 | $\beta$ | $\beta(1 - r_v)$ | $\beta(1 - r_f)$ |

In our model, the endemic pathogen has two genotypes: avirulent (*Avr*), which is sensitive to both forms of resistance, and virulent (*vir*) which is only sensitive to general resistance. Note that we follow the conventions of the gene-for-gene notation and use the term “virulent” to mean the specific ability to cause disease on a specifically resistant host (Flor, 1956; Thompson & Burdon, 1992), rather than as the amount of damage caused to the host, as it is more commonly used in the animal literature.

### Resistance Costs and Benefits

The baseline host-pathogen transmission rate is given by *β*, which can then be reduced by host resistance or costs of virulence (Table 1). We denote the resistance conferred by general resistance by *r*_*G*_ and the resistance conferred by specific resistance by *r*_*S*_. We also include each host genotypes’ susceptibility to a foreign pathogen. Following spillovers, foreign pathogens are often poorly adapted to their hosts (Lee et al., 2017). We therefore reduce the foreign pathogen’s transmission rate from the baseline by *r*_*f*_, for all host genotypes, which can be further reduced by general resistance.

Each form of resistance carries a cost to hosts in the form of a reduction in fecundity (*c*_*G*_ and *c*_*S*_). The resistance benefits of general and specific resistance, as well as their costs interact multiplicatively. The virulent genotype incurs a cost of virulence, given by *r*_*v*_, which reduces transmission on hosts lacking specific resistance (we notate this cost as *r*_*v*_ rather than *c*_*v*_ since it affects the pathogen’s transmission rate rather than host’s fecundity). This framework is based on Leonard’s “soft selection” gene-for-gene model (Leonard, 1994).

### SI Dynamics

We define the transmission matrix ***B*** by ***B***_*ij*_ = *β*_*ij*_ where *β*_*ij*_ is the transmission rate of the *j*th pathogen to the *i*th host (Table 1). Similarly, we define the resistance cost vector ***c*** by letting *c*_*i*_ be the resistance costs of the *i*th host (Table 1). Incorporating these assumptions into a frequency dependent SI model (Ross, 1916), we have the following system of ordinary differential equations.

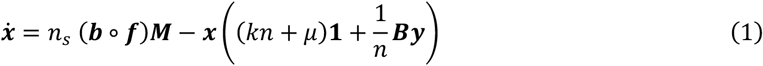

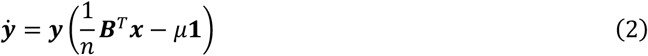

Here, ***x*** is the vector of uninfected host genotype abundances, and ***B*** is the vector of infected host abundances (for each pathogen genotype). The host population size is given by *n* (*n*_*s*_ is the total number of uninfected hosts). The vector **1** is a vector where every element is 1. The parameters *k* and *μ* represent the coefficient of host density-dependent growth and death rate, respectively. For uninfected hosts, new births occur via random mating, given by *n*_*s*_ (***b*** ∘ ***f***)***M***, where ***M*** is a two-locus mating matrix (later expanded to three locus, see evolution of linkage below) with recombination rate *ρ* (see below), ***b*** is the vector of maternal birthrates for each parental pair, and ***f*** is the probability of each parental pair (elementwise vector multiplication is denoted by ∘).

### Mating Matrix

Let *G* denote the genotype matrix, where *G*_*ij*_ ∈ {0,1} denotes the *j*th allele of the *i*th genotype. We assume that the loci are sequentially ordered, giving a total of 2^*n*^ genotypes, where *n* is the number of loci. We must then calculate the probability of maternal genotype *i*, and paternal genotype *j* producing offspring genotype *k* for all *i, j, k* ∈ {1, …, 2^*n*^}. To do so, we calculate the probability that an offspring will receive their allele at each locus from a particular parent.

Let *P*(*M*_*i*_) be the probability that an offspring receives their *i*th allele from their mother and *P*(*F*_*i*_) be the probability that their *i*th allele is from their father. We assume that *P*(*M*_1_) = *P*(*F*_1_) = 0.5. Next, let *ρ*_*i*_, *i* ∈ {1, …, *n* − 1} be the recombination rate between locus *i* and *i* + 1. We can then calculate the following allele origin probabilities inductively.

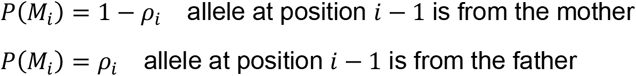

For each locus, the paternal origin probabilities are then given by *P*(*F*_*i*_) = 1 − *P*(*M*_*i*_). Since we assume recombination events are independent, the probability of the offspring’s genotypes coming from a particular sequence is the product of the probability of each allele. For any maternal, paternal, offspring combination, *i, j, k*, a subset of parental locus combinations will yield offspring genotype *k*. As these events are mutually exclusive, the probability of offspring *k* given maternal genotype *i* and paternal genotype *j* is the sum of the probabilities of each of these combinations. We then collapse the resulting 2^*n*^ × 2^*n*^ × 2^*n*^ tensor to a matrix with dimensions 2(2^*n*^) × 2^*n*^, such that the first index represents the parental genotype combination and the second index the offspring genotype.

### Transitivity Slope

For each set of parameters, we calculated the transitivity slope, based on the family-level susceptibilities to the foreign and endemic pathogens. With four host genotypes there are 16 fullsib families, given by the paternal-maternal genotype combinations. For full-sib family *i*, the family-level susceptibility to pathogen *j* is the average susceptibility off their offspring to pathogen *j*, weighted by the probability of the parental genotypes producing each offspring genotype.

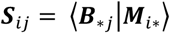

Here, ***B***_∗*j*_ is the transmission rate of pathogen *j* across each host genotype, and **M**_*i*∗_ is the vector of offspring probabilities for parental pair *i*, given by the mating matrix. This is also given by ***S*** = ***BM***. We consider the susceptibility to the endemic pathogen be the average across the *Avr* and *vir* genotypes, weighted by the relative abundance of each pathogen genotype. The transitivity slope is then computed by taking a weighted linear regression of all families’ average susceptibility to the endemic versus the foreign pathogen, weighted by likelihood of each family, ***f***. For simulations which yielded only a single type of full-sib family, we considered the transitivity to be zero.

### Numerical Analysis

Due to the high dimensionality and nonlinearity of our system, we were unable to find analytic expressions for our equilibria. We therefore used the modified Powell method for numerical root finding to estimate equilibrium points for our system, implemented with the Scipy 1.10.1 package in Python 3.11.2. (Powell, 1964; Virtanen et al., 2020). To ensure these estimated equilibria converged to interior (non-disease-free) equilibria, we first used the explicit Runge-Kutta method to compute numerical solutions, implemented with the SciPy 1.10.1. Each simulation was run to *t* = 5000. We then used the final point of these simulations as an initial guess for numerical root finding. To test the stability of these equilibria, we computed finite difference approximations of the Jacobian matrix at equilibria. We also computed numerical solutions to t = 10,000 and checked whether the solution at the endpoints were consistent with our numerical root finding (See Supplementary Material).

### Parameter Space Investigated

Throughout our investigation, we assumed that general resistance was weaker than specific resistance (*r*_*G*_ = 0.3, *r*_*S*_ = 0.9), as is often the case for plant pathogens (Poland et al., 2009). All other parameters were initially fixed as: *k* = 0.001,*μ* = 0.2, *b* = 1, *ρ* = 0.05, *β* = 0.5, *r*_*f*_ = 0.1. We began all simulations with the same initial conditions: 100 uninfected individuals of each of the four host genotypes and 10 hosts infected with the avirulent endemic pathogen genotype. For simulations with coevolution, we began with an additional 1 host infected with the virulent endemic pathogen genotype. We ran three series of simulations. First, we only included the avirulent endemic pathogen genotype to establish baseline equilibria without coevolution. In these simulations, we varied the cost of general resistance, and the cost of specific resistance (0 ≤ *c*_*G*_≤ 0.2, 0 ≤ *c*_*S*_ ≤ 0.4, values outside this range generally resulted in fixation at both loci). Next, we ran coevolutionary simulations including both endemic pathogen genotypes, also varying the costs of each form of resistance. Finally, we ran a set of coevolutionary simulations where we varied the cost of virulence instead of the cost of specific resistance (0 ≤*c*_*G*_ ≤ 0.2, 0 ≤ *r*_*v*_ ≤ 0.3, *c*_*S*_ was fixed at 0.2). For this last set of simulations, we opted to vary the cost of general resistance rather than the cost of specific resistance because the cost of general resistance had a more pronounced effect on evolutionary outcomes (see results). If polymorphism was maintained at both resistance loci, we calculated the degree of linkage disequilibrium (using Lewontin’s *D*^′^, see Supplementary Material), and the transitivity slope.

To assess the generality of our results, we ran a series of simulations with alternative assumptions. First, we investigated the impact of using density dependent transmission, with a baseline transmission rate of *β* = 0.001 to account for the different units of density-dependent transmission. We then tested the effects of the hard selection gene-for-gene model (Leonard, 1977), where the costs of virulence for the virulent pathogen genotype are applied equally against all host genotypes. Next, we investigated the effects of stronger general resistance, setting the strength of resistance, *r*_*G*_ to 0.5 (opposed to our baseline of 0.3). Finally, we tested the effects of the recombination rate, testing both full recombination *ρ* = 0.5 and complete linkage, *ρ* = 0.

### Evolution of linkage

Our results showed that the equilibria conditions were impacted by the degree of recombination, with the special case of complete linkage (*ρ* = 0) leading to markedly different outcomes than *ρ* = 0.05 (see Results). We therefore tested whether selection would favor the evolution of linked general and specific resistance. To do this, we added a third unlinked locus which modified the degree of linkage between *G* and *S*. We assume that this locus is unlinked from the general and specific resistance loci (*ρ*_1_ = 0.5). Each genotype’s recombination rate between general and specific resistance (*ρ*_2_) is then determined by the maternal linkage modifier allele. This framework allows us to model the ability for an allele which modifies recombination to invade. We then repeated the previous coevolutionary simulations with the additional linkage modifier locus, testing whether an allele which results in complete linkage (ρ_2_ = 0) between general and specific loci would evolve in a background of intermediate recombination (ρ_2_ = 0.05).

## Results

Our numerical approximation successfully converged for parameters examined. For nearly all parameters, the equilibria were asymptotically stable, per our numerical analysis. Exceptions to this generally occurred when costs of specific resistance or virulence were near zero and were likely due to numerical stability issues (see Supplementary Material, Fig S1-2). The results found though numerical root finding were qualitatively identical to our results found with numerical integration.

### The effect of coevolution on host evolution

Without coevolution, general and specific resistance were highly exclusionary, where hosts typically maintained only one form of resistance (contrast Fig 1A, D). Maintaining resistance at both loci was only possible when costs for both general and specific resistance were low (bottom left corner, Fig 1A, D). Allowing coevolution through the introduction of the virulent pathogen genotype expanded the range of conditions where general resistance was maintained (compare Fig 1A, B) and reduced the frequency of specific resistance (compare Fig 1D, E). This expansion of general resistance in response to coevolution was a robust result that occurred with density-dependent transmission (Fig S4A-C), alternative cost structuring for pathogen virulence (Fig S4D-F), as well as with both moderate (*r*_*G*_= 0.3) and high (*r*_*G*_= 0.5; Fig S4G-I) levels of general resistance.

**Figure 1.**
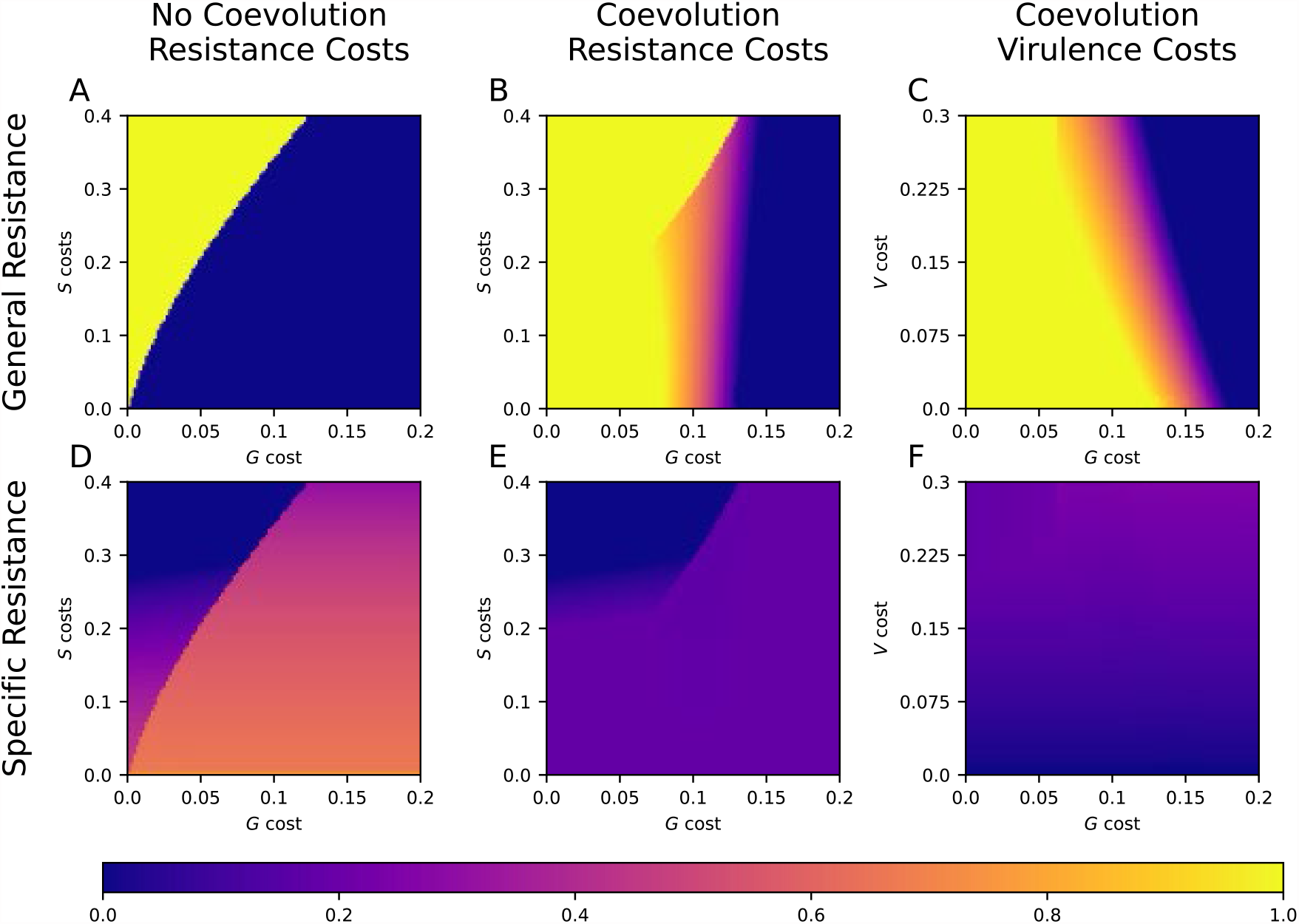
Host allele frequencies at equilibrium as a function of costs. Colors on the top row (A-C) show the frequency of the *G* allele, with warmer colors representing higher frequency. Colors on the bottom row (D-F) show the frequency of the *S* allele. (A, D): Frequency of the *G* and *S* allele across all hosts without coevolution as a function of *c*_*G*_ and *c*_*S*_. (B, E): Frequency of the *G* and *S* allele across all hosts with coevolution as a function of *c*_*G*_ and *c*_*S*_. (C, F): Frequency of the *G* and *S* allele across all hosts with coevolution as a function of *c*_*G*_and virulence costs (*r*_*v*_). Other parameters: *k* = 0.001, *μ* = 0.2, *b* = 1, *ρ* = 0.05, *β* = 0.5, *r*_*G*_= 0.3, *r*_*S*_ = 0.9.

Coevolution also expanded the range of conditions that led to stable polymorphism for general resistance. In our baseline scenario, when general resistance was moderate (*r*_*G*_= 0.3), polymorphism at the *G* locus could only be maintained with coevolution (orange region, Fig 1D). Higher levels of general resistance could drive the maintenance of *G*/*g* polymorphism in the absence of coevolution, but coevolution greatly expanded the region of stable polymorphisms (Fig S6). Coevolution led to the maintenance of simultaneous polymorphisms for general resistance (orange region, Fig 1B) and specific resistance locus (Fig 1E), enabling the maintenance of all four host alleles.

The frequencies of host resistance alleles depended on the costs of both general and specific resistance. Intermediate general resistance costs enabled polymorphism at both the general and specific resistance loci (orange zone, Fig 1B). However, costs of specific resistance did not have as strong an effect on the evolutionary outcomes. Very high costs of *S* (*c*_*S*_ > 0.25) could lead to the loss of specific resistance altogether (dark blue zone, Fig 1E) unless general resistance was also costly.

The cost of pathogen virulence influenced the equilibrium frequencies of both general and specific resistance (Fig 1C, F). Lower virulence costs increased the frequency of general resistance and decreased the frequency of specific resistance. With low virulence costs, the virulent genotype reached higher frequencies, and consequently selection for specific resistance was reduced since it was ineffective against the virulent genotype. As a result, the relative benefit of general resistance increased. Conversely, higher costs of virulence decreased the frequency of the virulent pathogen genotype, increased the frequency of *S*, decreased the frequency of *G*. Applying virulence costs to all host types did not strongly affect host allele frequencies (Fig S4D-F).

### The effects of recombination on general resistance

Increasing the recombination rate above our baseline *ρ* = 0.05 did not cause any substantial changes to allele frequencies (Fig S4J-L). The extreme case of complete linkage (*ρ* = 0, Fig 2), generated different outcomes, maintaining polymorphism in *G* even in the absence of coevolution (Fig 2A). In this case, the *Gs* and *gS* genotypes were maintained under frequency-dependent selection, while the *GS* and *gs* genotypes were lost. As a results, both general and specific resistance were polymorphic, even though only two genotypes were sustained. Coevolution expanded the region of polymorphism (orange area, Fig 2B), but as in the no coevolution scenario, only two genotypes (*Gs* and *gS*) were maintained.

**Figure 2.**
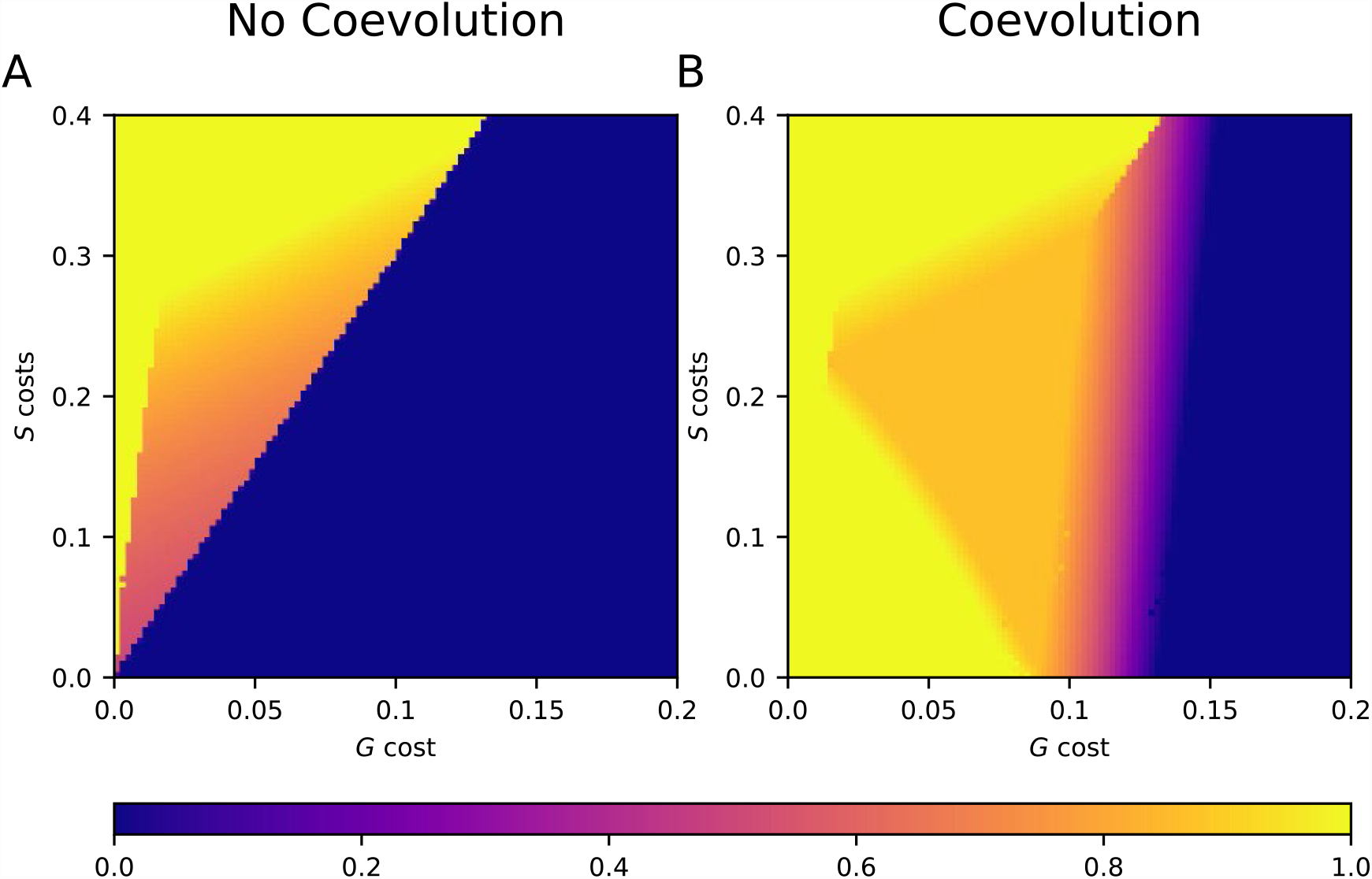
Frequency of the general resistance allele in simulations with no recombination. (A): Frequency of general resistance without coevolution. (B): Frequency of general resistance with coevolution. All other parameters besides the recombination rate (ρ = 0) are the same as in Fig 1.

### Determinants of resistance correlation

The cost of general resistance has a significant effect on the transitivity slope. If general resistance was lost or fixed, all genotypes had the same level of susceptibility to a foreign pathogen. Therefore, only parameters generating *G*/*g* polymorphism led to non-zero transitivity slopes. Since without coevolution, the *G* allele was either fixed or lost, coevolution was a requirement for non-zero transitivity slopes (assuming recombination). However, with intermediate *G* frequencies, there was a considerable region of positive transitivity (Fig 3).

**Figure 3.**
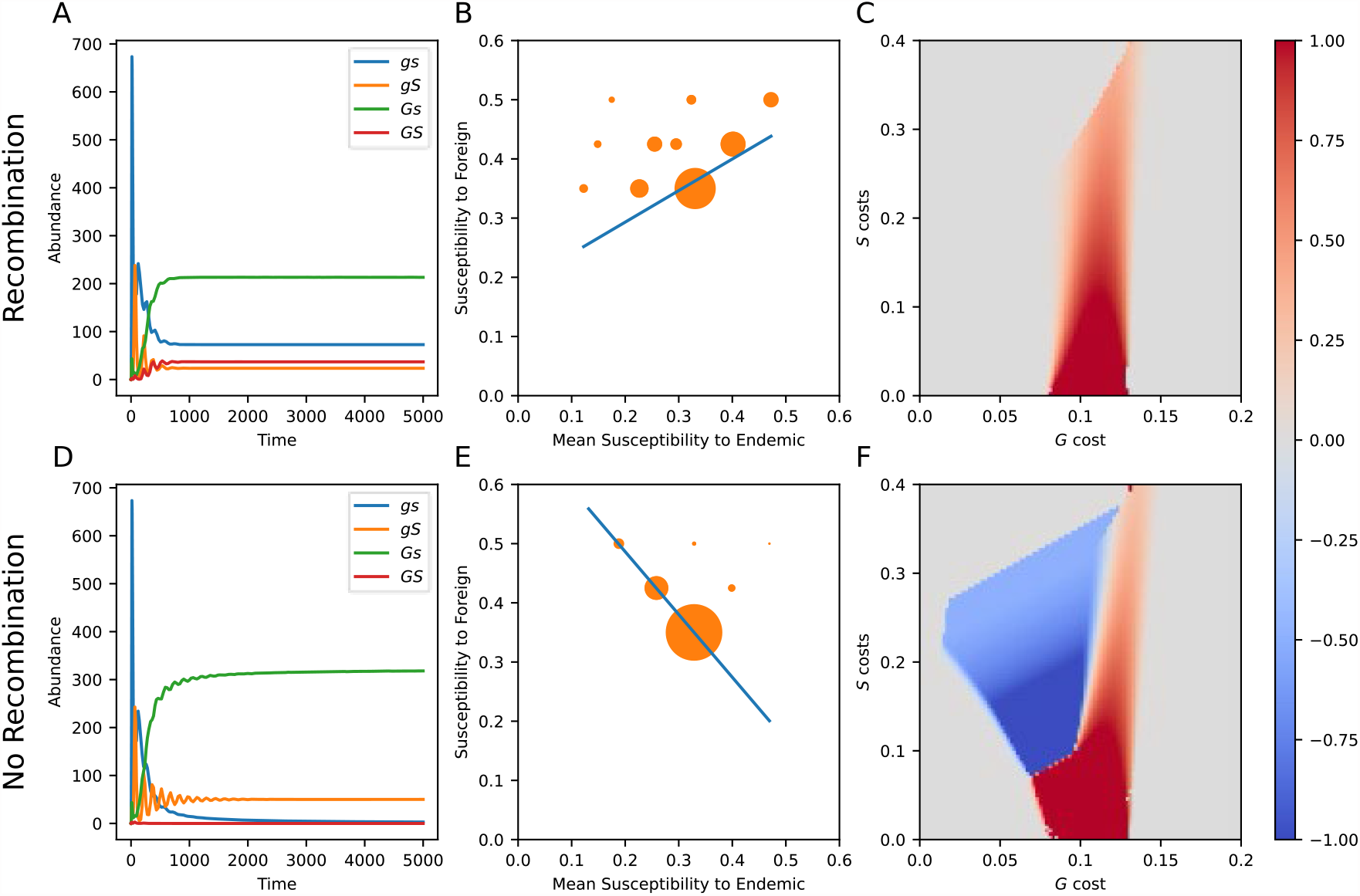
Resistance transitivity and the effects of the recombination rate. Panels A-C show results with the default recombination rate (*ρρ* = 0.05), while panels D-F assume no recombination (*ρ* = 0) (A, D): Host abundances for each genotype for (A) recombining and (D) non-recombining hosts. (B, E): Resistance transitivity across full-sib families for (C) recombining and (F) non-recombining hosts. The size of each symbol is proportional to the probability of randomly sampling each family. For A, B, D and E, *c*_*G*_= 0.1 and *c*_*S*_ = 0.2. (E, F): Transitivity slope for full-sib families with coevolution as a function of *c*_*G*_and *c*_*S*_. All parameters other than the recombination rate are the same as Fig 1.

Transitivity was highest when the frequency of *G* was at near 50%, and the cost of specific resistance was low (Fig 3C; see also Fig 1B, C for reference to the frequency of *G*). We found that the parameters most important to determining the slope of resistance transitivity were those that influenced the frequency of *G*, principally the cost of general resistance, *c*_*g*_ (Fig 3C). With low general resistance costs, *G* became fixed, and the transitivity slope was zero (as there is no variation in susceptibility to foreign pathogens). Generally, we found the highest transitivity values when the cost of specific resistance was low, however this may be less informative, as there was little between-family variation in susceptibility to the endemic pathogen.

A notable exception to the pattern of positive transitivity occurred when general and specific resistance were in complete linkage (*ρ* = 0, Fig 3D-F). With recombination rates above ∼0.02, and coevolution, *Gs* and *gs* were the most common genotypes (Fig 3A). However, without recombination, this switched to *Gs* and *gS* (Fig 3D), which were more fit without the loss of gametes to the costly double-resistant *GS* and highly susceptible *gs* genotypes. Given the high susceptibility to the foreign pathogen and low susceptibility to the endemic of *gS*, along with the even resistance of *Gs*, this resulted in negative transitivity (Fig 3E). The loss of recombination also led to a sharp boundary between positive and negative transitivity, where small changes in the cost of general resistance could significantly increase a populations susceptibility to spillover pathogens. Without recombination, positive transitivity is generated when *gs* and *Gs* become the most common genotypes (as opposed to *Gs* and *gS* for negative transitivity), driving an overall positively correlated trend.

### Evolution with a linage modifier allele

We found that a linkage modifier that eliminated recombination between general and specific resistance could invade and sweep to fixation in all cases where general resistance was polymorphic (Fig 4). In regions without general resistance polymorphism (also corresponding to regions with positive linage disequilibrium, see Fig S3), we found neutral selection on the two recombination alleles (white region, Fig 4C, Fig S2). However, if selected for, the time required for linkage modifier allele to go from introduction (50%) to fixation was significantly longer than it took for general and specific resistance to approach equilibrium (*t* ~ 5,000 vs *t* ~ 100,000, Fig 3A vs. Fig 4A) showing that while selection for tight linkage is often positive, it is also relatively weak. In many of these cases, the loss of recombination led a population which would otherwise have positive transitivity to switch to negative transitivity (Fig 4B).

**Figure 4.**
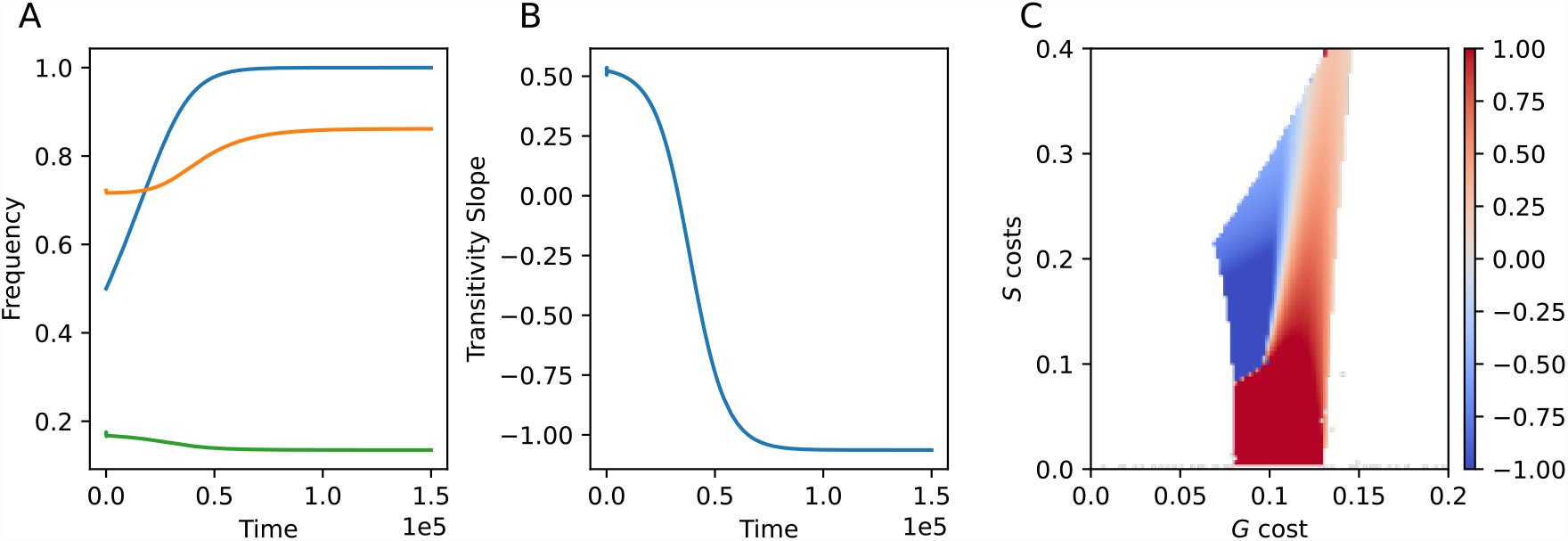
Allowing the evolution of recombination can result in complete linkage between general and specific resistance. Here, one linkage modifier allele causes a recombination rate of ρ = 0.05 while the other results in no recombination ρ = 0. (A): Allele frequencies for general resistance *G* (orange-line), and specific resistance *S* (green line), following the introduction of a linkage modifier allele (blue line). (B): Transitivity slope over time. (C): Parameters where the linkage modifier evolves to fixation, with colors showing the transitivity slope (as in Fig 3). White denotes regions where selection on the linkage modifier was neutral, owing to the lack of *G* polymorphism. All panels: parameters: *k* = 0.001, *μ* = 0.2, *b* = 1,*ρ* = 0.05, *β* = 0.5, *r*_*G*_= 0.3, *r*_*S*_ = 0.9. Panels A, B: *c*_*g*_ = 0.1, *c*_*s*_ = 0.2.

## Discussion

Emerging infectious diseases pose an increasing threat to agriculture and human health (El-Sayed & Kamel, 2020; Jones et al., 2008). While spillovers typically occur between phylogenetically related host species (Longdon et al., 2014), species often exhibit standing genetic variation for resistance to foreign pathogens (Gilbert et al., 2018; Marmor et al., 2006; McCarthy et al., 2011). This observation begs the question: what has led to the maintenance of this foreign pathogen resistance? Is it an evolutionary response to constant spillover events, or can it be explained as the outcome of coevolutionary dynamics with an endemic pathogen? Our results show that host-pathogen coevolution, in the absence of a foreign pathogen can drive the evolution of broad-spectrum, general resistance and the maintenance of genetic diversity within host populations.

In a previous study (Hulse et al., 2023), we found that under most conditions, evolution with an invariant endemic pathogen favored specific resistance while suppressing general resistance, maintaining a population’s susceptibility to foreign pathogens. However, our current results show that if an endemic pathogen can coevolve in response to specific resistance, general resistance is more often maintained, decreasing spillover risk. With coevolution, hosts maintained general resistance even when specific resistance had no costs, unlike in models without coevolution (Hulse et al., 2023). This is highly relevant to plant systems, where specific resistance costs have often been hard to detect (Brown, 2002; Korves & Bergelson, 2004). While our model was based on a frequency-dependent disease of wild plants, our key finding that coevolution expands the conditions for the maintenance of general resistance was robust to variation in transmission mode as we found similar outcomes with density-dependent diseases. This result was also robust across varying levels of general resistance and different forms of pathogen virulence costs.

A key prediction from our model is that gene-for-gene coevolution with specific resistance can drive the evolution of general resistance. The best way to test our predictions would be through either experimental evolution or time-shift experiments in natural populations, as both these approaches lead to direct estimates of dynamics. In a time-shift experiment with naturally occurring bacteria and phage inside leaves, Koskella and Parr (2015) found that while the phage rapidly adapted over time to track their local bacteria (suggestive of specific infectivity), the bacterial hosts evolved a more general form of resistance over time. Similarly, Fricken et al. (2016) found that experimental coevolution of algae and virus led to the evolution of general resistance in the algae. These results would seem to support our model, showing that general resistance is favored in the face of rapid coevolution. However, more recently Lewis et al. (2022), found the opposite trend, where *Caenorhabditis elegans* coevolved with a bacterial pathogen were more susceptible to foreign strains than the control hosts. In all cases however, the underlying infection genetic system is not known, making it challenging to draw direct comparisons with theory.

We also found that coevolution can lead to stable polymorphism for general resistance. In non-coevolving populations, moderate to low resistance is either fixed or lost, and polymorphism can only be maintained with high levels of resistance via ecological feedbacks (Antonovics & Thrall, 1994). Yet, we found that across all levels of general resistance, the addition of a coevolving pathogen expanded the region of polymorphism, and lead to greater genotypic diversity. This occurred even though coevolution occurred at the specific resistance locus, not the general resistance locus. Our results could therefore help explain how populations maintain genetic variation for foreign pathogens for which they do not share a coevolutionary history (Laine, 2004; Thrall et al., 2001).

Non-zero resistance transitivity never occurred unless there was also general resistance polymorphism. The success of foreign pathogen invasions can be affected by both the average level of foreign pathogen resistance as well as the slope of the resistance transitivity, especially with negative transitivity (Lerner et al., 2021). Maintaining individuals in a population that are more susceptible to foreign pathogens than to endemic pathogens creates an ecological niche for foreign pathogens to invade and persist. In our model, with even modest recombination, we only observed positive transitivity. This result recapitulates the results found by Lerner et al. (2021) for *Silene vulgaris*. However, with complete linkage resistance transitivity could become strongly negative because hosts either carried only the specific resistance gene or general resistance gene, but rarely both. Furthermore, we found that when the recombination rate is allowed to evolve, selection is neutral or favors complete linkage. This result is synergistic with empirical evidence, as in plants, resistance genes can be closely linked to begin with. In many species, *R* genes are clustered together in the genome (Hulbert et al., 2001; Meyers et al., 2005), and are thought to have evolved through tandem gene duplication (Couch et al., 2006; Mcdowell & Simon, 2006).

In our model, general resistance was conferred by a single gene that conveyed moderate levels of broad-spectrum resistance. While general resistance is more typically considered to be a polygenic trait, with many loci of small effect (Poland et al., 2009), there is considerable value in understanding the evolutionary dynamics at a single general resistance locus, before adding additional complexity. Indeed, our results disrupt a common view of general resistance as evolutionarily “durable”, and unaffected by pathogen coevolution (Brown, 2015). Moreover, major-gene general resistance has been described in plants, for example, the *PigmR* gene in rice is a typical nucleotide-binding, leucine-rich repeat (NLR) gene that protects against a large breadth of *Magnaporthe orzye* genotypes (Deng et al., 2017). In addition, high-temperature adult resistance (HTAP) genes in wheat have been shown to confer moderate levels of resistance against a broad-spectrum of leaf rust strains (Chen, 2013). However, future expansion of the model to explicitly consider a quantitative form of general resistance could provide new insights, particularly regarding the evolution of linkage.

Our coevolutionary model for specific resistance was based on gene-for-gene interactions, which are common in plant-pathogen systems (Thompson & Burdon, 1992). It is possible that a different infection genetics, such as matching alleles (Grosberg & Hart, 2000), could alter the outcome. In a matching alleles model, the ability to infect one host genotype comes at cost of the complete inability to infect other host genotypes. Polymorphism in both hosts and pathogens can be maintained without costs due to strong negative frequency-dependent fluctuating selection (Agrawal & Lively, 2002). It therefore seems unlikely that costly general resistance would be favored with matching alleles. In contrast, gene-for-gene interactions require both costs of resistance and virulence for the maintenance of polymorphism (Tellier & Brown, 2007). Our model shows that this system can be invaded by general resistance when either the costs of that general resistance is low, or the cost of pathogen virulence is low (rendering specific resistance less useful). In the future, expansion of the model to multi-locus gene-for-gene dynamics (Sasaki, 2000), or intermediate infection genetic models (Boots et al., 2014) could provide additional insights.

Our results demonstrate how pairwise coevolution between hosts and pathogens at a single resistance locus can ripple out and affect evolutionary dynamics at other resistance loci via feedbacks between host and pathogen, with broader consequences for cross-species transmission. Previous research has demonstrated the importance of feedback between general and specific resistance (Frank, 2000; Hulse et al., 2023), although less is known about how these dynamics are affected by coevolution. Coevolutionary arms-race dynamics have been shown in many systems (Grenfell et al., 1995), such as the gene-for-gene system in plants (Flor, 1956), with major implications for natural populations as well as agricultural systems. However, the focus has mostly been on the evolution of specific resistance genes themselves. By incorporating the evolution of general resistance into a coevolutionary model, our results demonstrate how coevolving endemic pathogens can make host populations more resilient to spillover.

## Author Contributions

S.V.H., J.A., M.E.H., and E.L.B. designed research; S.V.H., and E.L.B. performed research; S.V.H., and E.L.B. analyzed data; S.V.H. and E.L.B. wrote the paper with input from J.A., and M.E.H. All authors agreed on the final manuscript.

## Acknowledgments

This work was supported by a grant from the NIH (grant number R01GM140457) to Michael Hood and Emily Bruns.

## Data Availability

All code used to generate our data is available via GitHub at https://github.com/svhulse/gfg-model.

