## Supplementary Materials for "Host-pathogen coevolution promotes the evolution of general, broad-spectrum resistance and reduces foreign pathogen spillover risk"

### Linkage Disequilibrium Calculation

We calculated linkage disequilibrium (LD) based on the equilibrium allele frequencies (Eq S1).

$$D = f(GS) - f(G)f(S) \qquad (S1)$$

We then calculated $D'$ to normalize LD based on the maximum possible LD given the allele frequencies (Eq S2).

$$D' = \frac{D}{D_{max}} \qquad (S2)$$

Where

$$D_{max} = \max\{-f(G)f(S), -(1-f(G))(1-f(S))\} \text{ when } D < 0$$
$$D_{max} = \min\{f(G)(1-f(S)), (1-f(G)f(S))\} \text{ when } D > 0$$

Additionally, we calculated the LD for both the no recombination and full recombination simulations (Fig S1-2).

### Numerical Integration Results

To verify that our numerical root finding was converging to equilibria that would be attained in biologically feasible scenarios, we ran simulations using the same parameters as in the main text but running numerical integration to $t = 10{,}000$. To account for solutions with oscillatory equilibria (or very gradually dampened oscillations), we averaged each solution over the last 2000 time point to obtain an approximate equilibrium. We found no major differences in outcomes using this method when compared to the methods described in the main text.

### Stability Analysis Results

With our coevolutionary simulations, nearly all parameters tested produced asymptotically stable equilibria (Fig S2). Cases where equilibria had a non-negative eigenvalue corresponded either to parameters near boundaries of genotype fixation or loss or regions where gene-for-gene coevolution can produce highly irregular oscillatory dynamics. In either of these cases, the non-stability of equilibria is likely due to numerical issues, and these scenarios are rare enough to not influence our conclusions.

We also assessed the equilibrium stability for our three-locus model with the linkage modifier allele (Fig S3). We examined equilibrium stability when the modifier locus was either fixed or lost. With loss, there was a significant region where the equilibria were unstable, but the same parameters corresponded to a stable equilibrium when the linkage modifier was fixed. These parameters represent scenarios where if the modifier allele can invade, it will sweep to fixation, potentially shifting a population from positive to negative resistance transitivity.

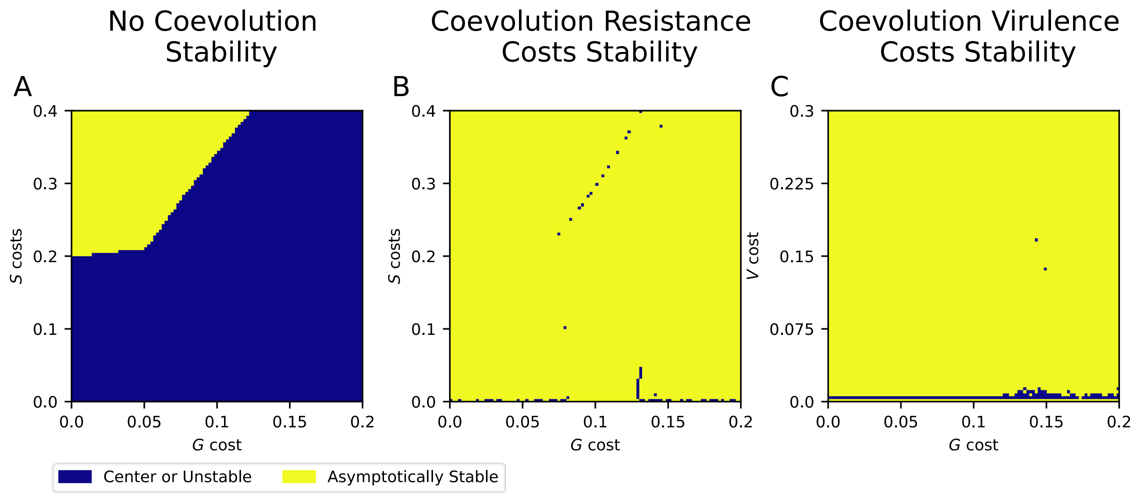


**Figure S1. Equilibria classifications for default parameters.** Yellow colors denote asymptotically stable solutions while blue represent solutions with a non-negative eigenvalue. (A): Stability for solutions without coevolution. Here, the large area of unstable solutions corresponds to regions where introducing the virulent pathogen genotype changes the equilibrium solution. (B): Stability for solutions with coevolution as a function of $c_g$ and $c_s$. (C): Stability for solutions with coevolution as a function of $c_g$ and virulence costs ($\boldsymbol{r_v}$).

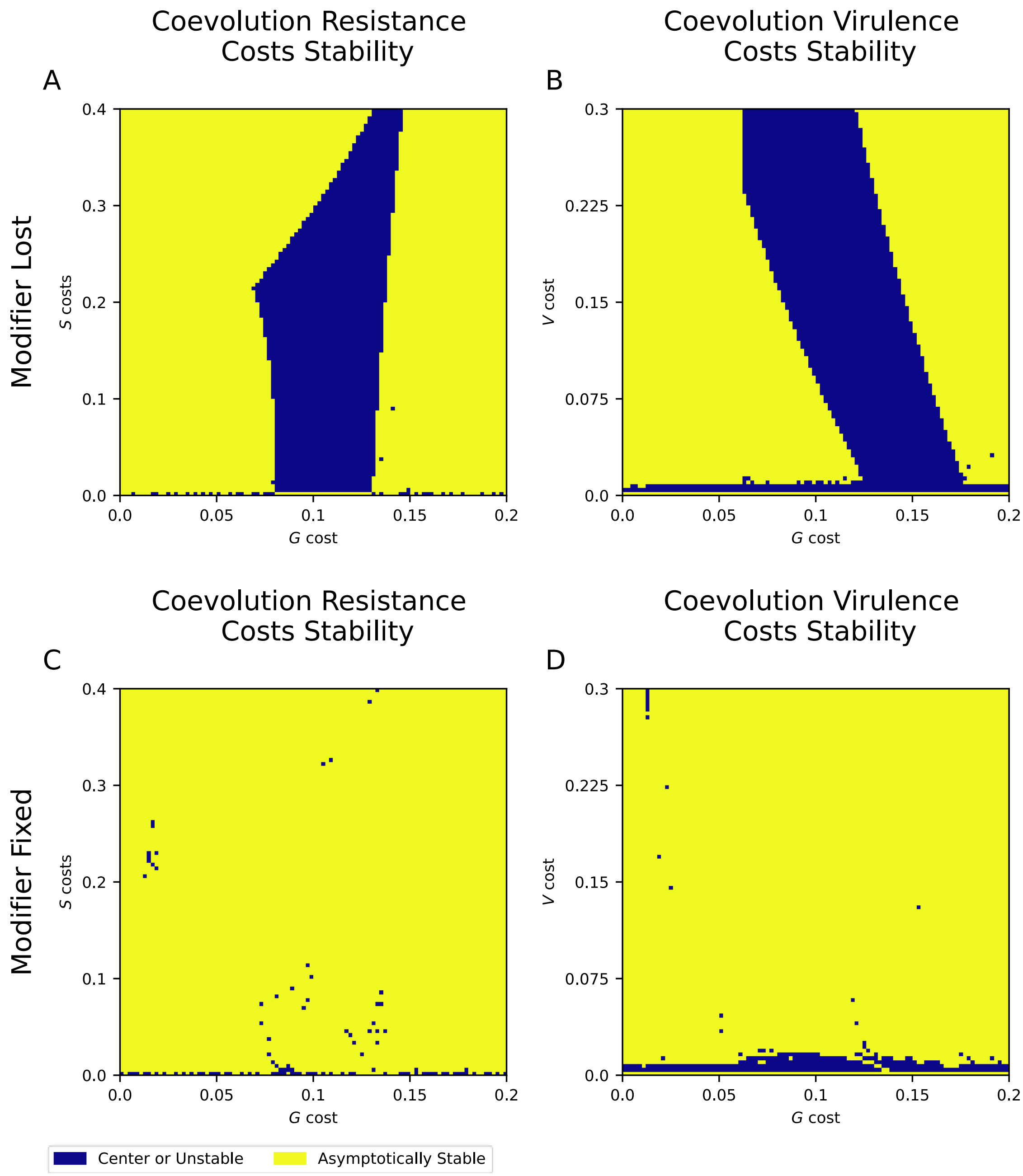


**Figure S2. Equilibria classifications for solutions in the three-locus model.** Yellow colors denote asymptotically stable solutions while blue represent solutions with a non-negative eigenvalue. (A-B): Stability for solutions where the modifier locus is lost as a function of (A): $c_g$ and $c_s$ and (B): $c_g$ and $r_v$. Blue regions indicate parameters where the modifier locus can invade and evolve to fixation. (C-D): Stability for solutions where the modifier locus is fixed as a function of (C): $c_g$ and $c_s$ and (D): $c_g$ and $r_v$.

### Linkage Disequilibrium

If polymorphism was maintained at both resistance loci, we calculated the degree of linkage disequilibrium (LD), and the transitivity slope. For LD, we used Lewontin's $D'$ normalized LD measure, where a value of 1 represents the highest possible level of linkage disequilibrium for any given allele frequencies. We also recorded the sign of LD, as this tells us the relative frequency of the *GS* genotype based on allele frequencies. To avoid numerical issues associated with low allele frequencies, we only considered LD for simulations where each allele represented at least 1 percent of the uninfected host population.

We found non-zero linkage disequilibrium (LD) across parameters where general resistance was polymorphic (Fig S2). When LD was non-zero, it was always negative, indicating a lower relative fitness of the *GS* and *gs* genotypes. Without coevolution, general resistance was either fixed or lost, there were no parameters which led to non-zero LD.

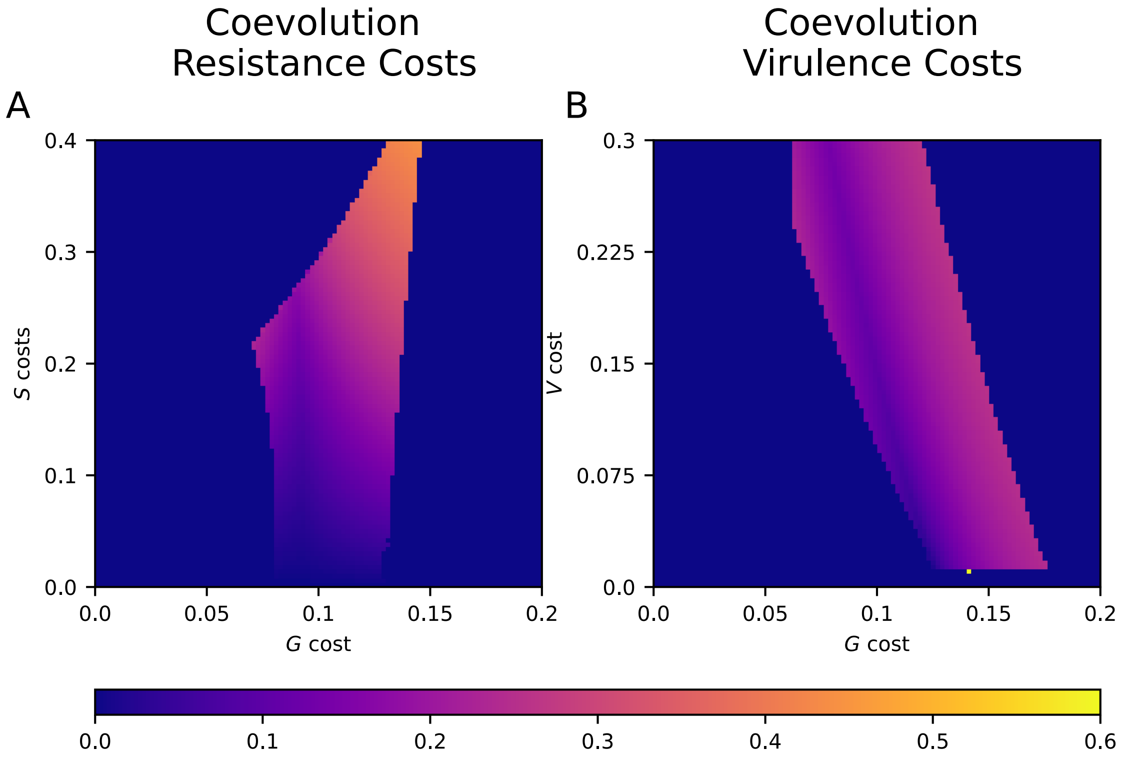


**Figure S3. Normalized linkage disequilibrium.** Normalized linkage disequibrium ($D'$) is shown as a function of (A) general and specific resistance costs and (B) general resistance and virulence costs. A D' value of 1 represents the highest possible LD given the equilibrium allele frequencies. All other parameters are the same as Fig 1.

### Alternative Model Assumption Results

We considered a range of alternative models, using density-dependent transmission (Fig S4), as well as the hard selection gene-for-gene model (Fig S3D-F), where virulence costs apply to all host genotypes. The equations for our system with density-dependent transmission are given below (Eq S3-4).

$$\frac{d\boldsymbol{x}}{dt} = n_s\,(\boldsymbol{b} \circ \boldsymbol{f})\boldsymbol{M} - \boldsymbol{x}\big((kn + \mu)\mathbf{1} + \boldsymbol{B}\boldsymbol{y}\big) \qquad (S3)$$

$$\frac{d\boldsymbol{y}}{dt} = \boldsymbol{y}(\boldsymbol{B}^T\boldsymbol{x} - \mu\mathbf{1}) \qquad (S4)$$

Additionally, we tested alternative default parameters, including stronger general resistance (Fig S4G-I), and variable recombination rates (Fig S4J-L).

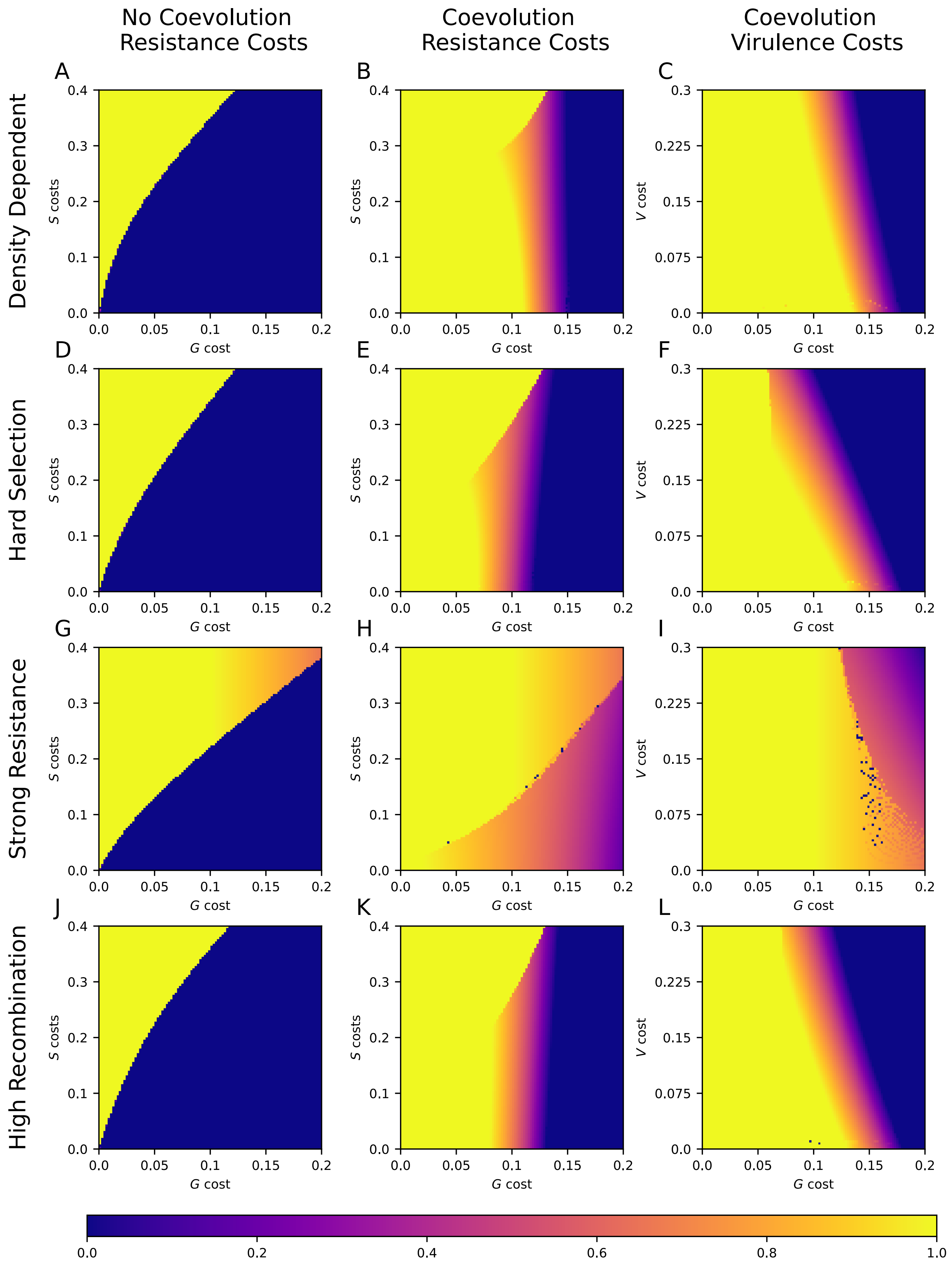

No Coevolution Resistance Costs
Coevolution Resistance Costs
Coevolution Virulence Costs
Density Dependent
Hard Selection
Strong Resistance
High Recombination
A
B
C
D
E
F
G
H
I
J
K
L
S costs
V cost
G cost
0.0
0.1
0.2
0.3
0.4
0.05
0.15
0.075
0.225
0.6
0.8
1.0

**Fig S4. General resistance allele frequencies in alternative models.** (A-C): Density-dependent transmission, with $\beta = 0.001$ throughout. (D-F): Hard-selection gene-for-gene model, where the costs of virulence apply to all host genotypes. (G-I): Model with stronger general resistance ($r_G = 0.5$). (J-L): Model with full recombination ($\rho = 0.5$). Unless stated, all other parameters are the same as in Fig 1.

**Table S1.** Mating matrix example for the two-locus model showing the genotype frequencies for offspring of each possible parental pair. To calculate the total proportion of offspring genotypes, these frequencies are weighted by the frequency of each parental pair, assuming random mating. The matrix $\boldsymbol{M}$ in the main text is given by the last four columns here.

| Parental Genotype 1 | Parental Genotype 2 | $GS$ Offspring Frequency | $Gs$ Offspring Frequency | $gS$ Offspring Frequency | $gs$ Offspring Frequency |
|---|---|---|---|---|---|
| $GS$ | $GS$ | $1$ | $0$ | $0$ | $0$ |
| $GS$ | $Gs$ | $0.5$ | $0.5$ | $0$ | $0$ |
| $GS$ | $gS$ | $0.5$ | $0$ | $0.5$ | $0$ |
| $GS$ | $gs$ | $0.5(1-\rho)$ | $0.5\rho$ | $0.5\rho$ | $0.5(1-\rho)$ |
| $Gs$ | $GS$ | $0.5$ | $0.5$ | $0$ | $0$ |
| $Gs$ | $Gs$ | $0$ | $1$ | $0$ | $0$ |
| $Gs$ | $gS$ | $0.5\rho$ | $0.5(1-\rho)$ | $0.5(1-\rho)$ | $0.5\rho$ |
| $Gs$ | $gs$ | $0$ | $0.5$ | $0$ | $0.5$ |
| $gS$ | $GS$ | $0.5$ | $0$ | $0.5$ | $0$ |
| $gS$ | $Gs$ | $0.5\ \rho$ | $0.5(1-\rho)$ | $0.5(1-\rho)$ | $0.5\ \rho$ |
| $gS$ | $gS$ | $0$ | $0$ | $1$ | $0$ |
| $gS$ | $gs$ | $0$ | $0$ | $0.5$ | $0.5$ |
| $gs$ | $GS$ | $0.5(1-\ \rho)$ | $0.5\ \rho$ | $0.5\ \rho$ | $0.5(1-\ \rho)$ |
| $gs$ | $Gs$ | $0$ | $0.5$ | $0$ | $0.5$ |
| $gs$ | $gS$ | $0$ | $0$ | $0.5$ | $0.5$ |
| $Gs$ | $gs$ | $0$ | $0$ | $0$ | $1$ |
